# HAPLODIPLOIDY ACCELERATES MITOGENOME EVOLUTION IN INSECTS

**DOI:** 10.1101/2024.12.02.626394

**Authors:** Avas Pakrashi, Paul DN Hebert

## Abstract

Rates of mitogenome evolution differ among animal lineages, variation which has been linked to life history, ecological traits, and potentially to breeding system. Insects are a good model for examining the latter impacts as, although most are diplodiploids (DD), some lineages reproduce by haplodiploidy (HD) or thelytoky. In this study, we ask if breeding system influences patterns of evolution in the mitogenome using the 658 bp barcode region of the cytochrome *c* oxidase I (COI) gene as a sentinel. Specifically, we ask if the incidence of amino acid substitutions and indels is linked to breeding system. We investigated this matter by examining COI sequences from specimens assigned to 1332 BINs, a species proxy. Belonging to 513 families and 26 orders, these BINs included representatives of roughly half of all insect families. Most are DD, but ten lineages, varying in rank from tribe to order, are HD. Our analysis reveals that HD lineages show higher rates of amino acid substitution than DD. In addition, indels are frequent in HD lineages but are absent in DD taxa with one exception. Among DD, the Strepsiptera is an outlier; its COI shows very rapid amino acid evolution and frequent indels. The accelerated rates of mitogenome evolution in HD lineages suggest that this breeding system facilitates coevolution between the nuclear and mitochondrial genomes.

## Introduction

Rates of molecular evolution vary among taxonomic lineages and across genomic compartments. For example, protein-coding genes in animal mitochondria evolve approximately 10 times faster than their nuclear counterparts (Brown et al., 1979). However, rates of mitogenome evolution also vary among animal lineages (Denver et al., 2000; James et al., 2016). Prior investigations have linked this variation to flight loss, metabolic rate, body mass, generation length, and parasitic lifestyle (Gillooly et al., 2005; Mitterboeck et al., 2013; Nabholz et al., 2008; Lynch et al., 2010, McMahon et al., 2011). However, the explanation for one one of the most striking differences in mitogenome evolutionary rates has lacked explanation. Early phylogenetic analyses revealed very accelerated mitogenome evolution in the honeybee (Crozier et al., 1977). While this was initially attributed to eusociality, subsequent studies revealed similar acceleration in solitary hymenopterans (Crozier et al., 1989). The hypothesis that HD, an uncommon breeding system shared by all Hymenoptera, explained their accelerated evolution was considered, but dismissed due to the difficulty in understanding how male haploidy could influence evolution of the maternally inherited mitogenome (Dapper et al., 2022). However, the matter is not settled as Bendall et al. (2022) suggested that haplodiploidy can expedite genomic differentiation and even foster sympatric speciation under divergent selection. More recently, Li et al. (2024) provide evidence for the accelerated evolution of mitochondrion-related genes in a subset of haplodiploid arthropods.

Because HD (both arrhenotoky and paternal genome elimination) has independently evolved on at least six occasions in insects, it is possible to test the generality of rate acceleration in groups employing this mode of reproduction. All species in two insect orders (Hymenoptera, Thysanoptera) are HDs and so too are some lineages of Hemiptera, Psocodea, Diptera, and Coleoptera (De La Filia et al., 2015, Blackmon et al., 2017, Joshi et al., 2023). Within the Hemiptera, HD occurs in the Aleyrodidae (white flies) and in all 17 families of the Coccoidea (scale insects) (Gullan et al., 2007; Blackmon et al., 2017; Jaron et al., 2022). In the Psocodea, HD occurs in the Liposcelididae and in several parasitic lice families (Phthiraptera) (Hodson et al., 2017). Within the Diptera, HD is employed by all Cecidomyiidae and by some Sciaridae (Herbette et al., 2023). Finally, in the Coleoptera, HD occurs in the sole species of Micromalthidae and in some species in 2 of the 19 tribes (Cryphalini, Xyloborini) of Scolytinae, one of the 19 subfamilies of Curculionidae (Marvaldi et al., 2005; Kawasaki et al., 2016; Blackmon et al., 2017). Thelytoky, asexual reproduction, occurs in most insect orders (Vershinina et al., 2016; Gokhman et al., 2018, Aguín-Pombo et al., 2023) but because thelytokous lineages are short-lived as evidenced by the lack of genera or even monophyletic assemblages of sister species, we do not consider them further (Blackmon et al., 2017).

In this study, we consider patterns of sequence diversity in the 658 bp barcode region of the mitochondrial cytochrome *c* oxidase subunit I (COI) gene (Hebert et al., 2003) across the Insecta. We examine two aspects of sequence diversity in this gene region **–** the incidence of amino acid (aa) substitutions and of indels (insertions/deletions). We compare these traits across 513 families belonging to 26 insect orders. Aside from comparisons between the two purely HD (Hymenoptera, Thysanoptera) and 20 purely DD orders, we compare HD and DD lineages in the four orders (Coleoptera, Diptera, Hemiptera, Psocodea) with mixed breeding systems.

## Materials and Methods

### Dataset preparation

A dataset (DS-HDMSDAT2) was assembled with representatives from all insect families with over 500 publicly available records on BOLD (https://boldsystems.org/). A few BINs from each of these families were chosen based on two criteria: (1) the barcode sequence length was > 646 base pairs, and (2) a photograph was available for the specimen representing the BIN. For 279 families, only one or two BINs met these criteria, but 3-8 BINs were haphazardly selected from each of the remaining families excepting the Curculionidae. In this family, 72 BINs were selected to include the broadest possible representation of its subfamilies and tribes. This approach produced a dataset with a single representative of 1332 BINs belonging to 513 families and 26 orders (**Table S1**).

### Consensus amino acid dataset and substitution analysis

The 1332 sequences were aligned using MAFFT (Katoh et al., 2002) and translated into amino acids using the “invertebrate mitochondrial” code in MEGA11. A secondary dataset was then constructed with the consensus aa sequence for each of the 513 families, employing the “most common bases” criterion in Geneious Prime 2024. Using a similar protocol, a consensus aa sequence was determined for two orders (Diplura, Protura) of Entognatha, the subphylum most closely related to the Insecta to serve as reference ancestral sequences. The number of aa differences between the consensus sequence for each insect family and the two outgroup sequences was then calculated in Geneious Prime. Radar plots and boxplots of results were generated using Plotly (Sievert, 2020) and ggplot2 (Wickham, 2016) respectively within Rstudio (RStudio Team, 2020). The pairwise aa distance between the consensus sequence for each family in each order was also calculated using the Poisson model as a measure of amino acid variation among taxa in each order (Nei, 1987) in MEGA11 (Kumar et al., 2018).

### Mapping Indels

The occurrence and placement of insertions and deletions in the consensus aa sequence for each family was ascertained by comparison with the consensus sequence for all insects. The secondary structure of the consensus aa sequences was predicted using SWISS-MODEL (Waterhouse et al., 2018) and subsequently visualized in Jalview (Waterhouse et al., 2009). Helix and loop numbering followed Pentinsaari et al. (2016). The resultant figures were further edited using Inkscape 1.3.2.

### Phylogenetic analysis

Phylogenetic analysis of the consensus aa dataset for the 513 families utilized the Bayesian method implemented in BEAST2.6.3 (Bouckaert et al., 2014). The GTR+I+G substitution model, empirical base frequencies, and strict clock with the Yule model were employed as tree priors. To reduce exposure to long-branch attraction, the phylogeny was constrained to ensure order-level monophyly.

Two independent Markov Chain Monte Carlo (MCMC) runs were executed with each extending for 1,000,000 generations. The initial 25% of each run was discarded as burn-in, and sampling was conducted every 1000 generations. Subsequently, LogCombiner V1.5.3 was employed to merge results from the runs. The adequacy of mixing of the MCMC chains was assessed using Tracer V1.7.1 with the goal of ensuring that all parameters attained an effective sample size (ESS) > 200. Following removal of the first 10% of trees from the combined tree file, a maximum clade credibility tree was generated using TreeAnnotator 1.5.4 with a posterior probability limit set at 0.5 (Bouckaert et al., 2014). The resulting phylogenetic tree was visualized using FigTree 1.4.4 (Rambaut, 2018) and edited using iTOL (Letunic et al., 2021).

## Results

### Amino acid substitutions in insect COI

The barcode region of COI spans 220 aa and 177 of these positions showed one or more substitutions among the 1332 BINs. The number of aa substitutions between the dipluran consensus and the consensus sequences for the 513 insect families had a greater range (25**–**120) than with the proturan (51**–**114) (**Figure 1**). However, the number of substitutions between each insect family and the dipluran/proturan outgroups was strongly correlated (r^2^ = 0.82). Because of this congruence (**Table S2**), subsequent analyses employed values from the dipluran comparison (**Table S3**). The two subclasses and three superorders comprising the Insecta showed nearly 2-fold variation in their mean number of aa substitutions from the dipluran (Apterygota**–**33, Palaeoptera**–**33, Polyneoptera**–**39, Holometabola**–**42, Paraneoptera**–**54) (ANOVA, F= 18.2, p<0.001) (**Figures S1**-**S10**, **Table S4**).

**Figure 1:**
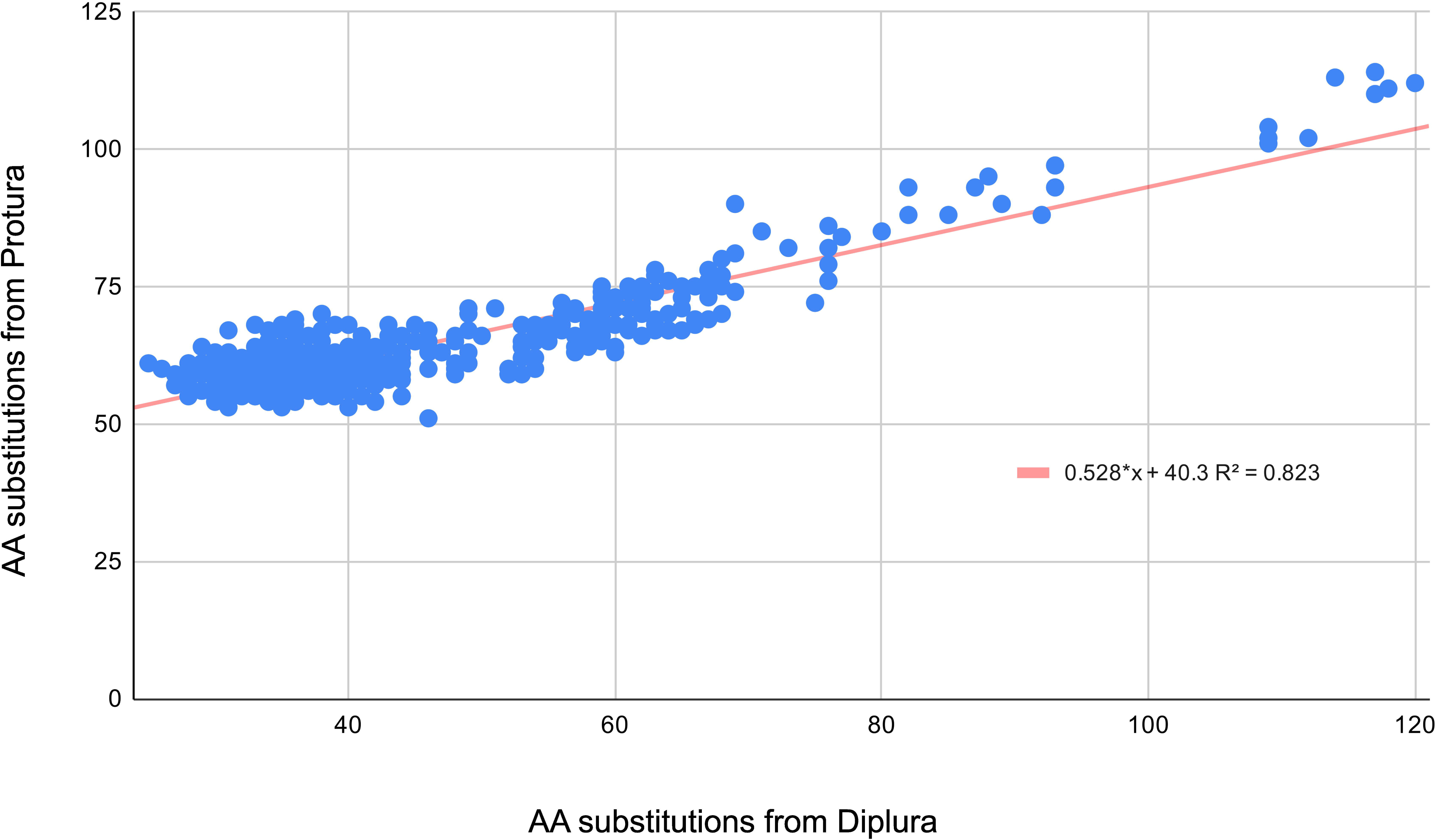
Relationship between the number of amino acid substitutions for a consensus dipluran sequence and a consensus proturan sequence and the consensus sequence for each of 513 insect families.

Considering all families, the average number of substitutions was 44 with 20% having >50 substitutions, 3% >75 substitutions, and 2% >100 substitutions. To aid subsequent discussion, we classed families in the upper quintile (>50 substitutions) as having a high number of substitutions while those in the lowest quintile (<34 substitutions) were classed as having few substitutions. The mean number of substitutions varied nearly 2-fold (ANOVA, F= 75.19, p<0.001) among families in the five major insect orders **–** from a low of 34 substitutions in Diptera to a high of 62 in Hymenoptera (**Figure 2A**, **Figure S1–S5**, **Table S5**). Considering all 26 insect orders, Strepsiptera (89) and Thysanoptera (66) had more substitutions than Hymenoptera (62) which was followed by Hemiptera (55), Embioptera (55), and Phasmatodea (54) (**Figure 2B**). Ten orders (Archeognatha**–**33, Zygentoma**–**33, Siphonoptera**–**32, Dermaptera**–**32, Mecoptera**–**31, Neuroptera**–**31, Ephemeroptera**–**30, Plecoptera**–**29, Raphidoptera**–**29, Megaloptera**–**27) joined the Diptera in the lowest quintile. Orders in the upper quintile also tended to show marked amino acid divergence among their component families as the five orders with the highest inter-family divergence were Strepsiptera (34), Hemiptera (31), Hymenoptera (28), Thysanoptera (23), and Embioptera (19) (**Figure 2B**, **Table S3**).

**Figure 2:**
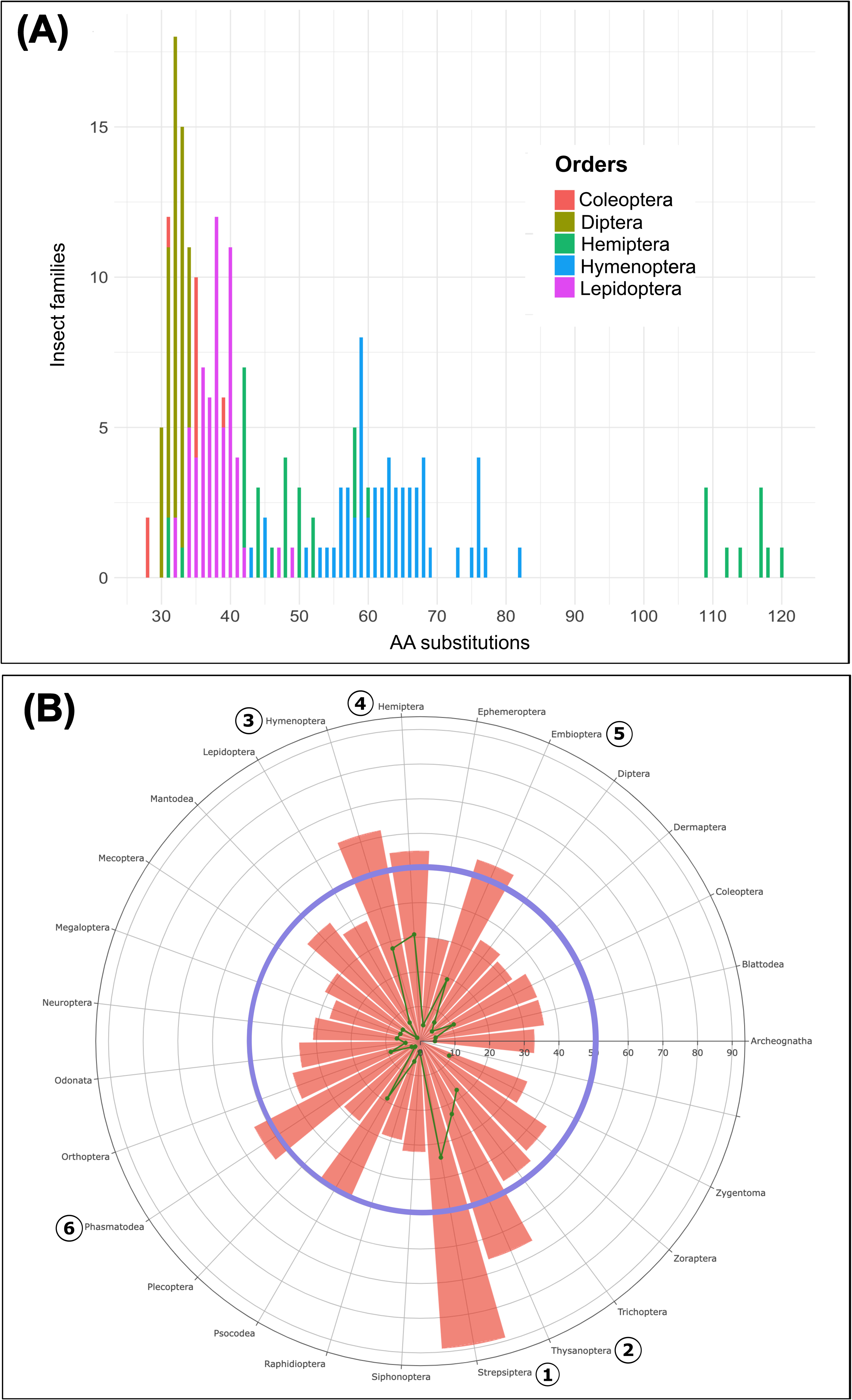
(A) Histogram of the number of amino acid substitutions between the consensus sequence for a dipluran outgroup and 368 families in the five major orders of insects. (B) Radar plot of the mean number of amino acid substitutions between a dipluran outgroup and 26 orders of insects. The blue annulus indicates 50 substitutions. Interfamily Poisson genetic distance showing in internal scatter green plot.

### Amino acid substitutions within orders with haplodiploid lineages

Only two insect orders reproduce solely by HD, and nearly all their component families fell in the upper quintile although substitution counts varied among their component lineages. For example, within the Thysanoptera, the sole family (Phlaeothripidae) in the suborder Tubulifera had more substitutions (71) than the three families (Heterothripidae**–**67, Thripidae**–**64, Aeolothripidae**–**63) in the suborder Terebrantia (**Figure 3**, **Figure S8**). Among the 62 families of Hymenoptera, 57 were in the upper quintile, but the 51 families in the suborder Apocrita had more substitutions than the 11 in the suborder Symphyta (64 vs 53; t = 2.9, p<0.05) (**Figure 3**, **Figure S4**, **Table S6**).

**Figure 3:**
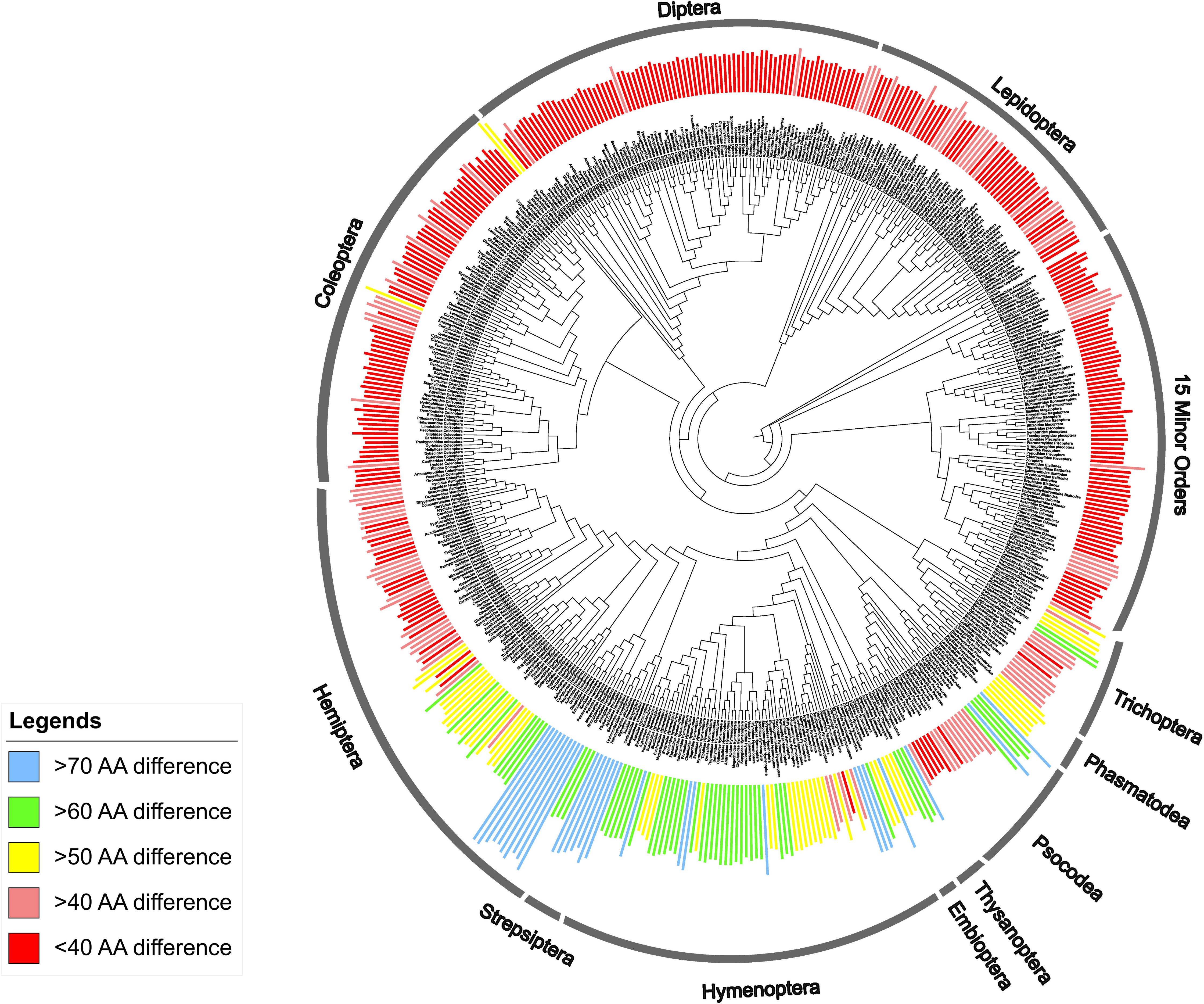
Order constrained Bayesian phylogeny for 513 families of insects. Colours show the number of amino acid substitutions between a dipluran outgroup and each of these families.

Among the four orders with mixed breeding systems, HD lineages regularly showed more aa substitutions than DD lineages.

#### Hemiptera

Considerable variation was apparent in the mean number of aa substitutions among the three suborders of Hemiptera (Heteroptera**–**40, Auchenorrhyncha**–**53, Sternorrhyncha**–**89) (**Figure S5, Table S7**). All 38 families of Heteroptera are DD and just one (Schizopteridae) fell in the upper quintile. All Auchenorrhyncha are also DD, but the seven families in the infraorder Cicadimorpha had fewer substitutions than the 17 families in the infraorder Fulgoromorpha (45 vs 56). The 19 families of Sternorrhyncha collectively employ both the breeding systems. The five DD families (superfamily Psylloidea) had comparatively low substitutions (mean 59)(**Table S7**). The 11 HD families were all in the upper quintile although the number of substitutions (63) for Aleyrodidae (whiteflies) was much lower than those for the 10 families of Coccoidea (x = 114) (**Figure 3**, **Figure S5**, **Table S7**).

#### Psocodea

Within the Psocodea, members of the suborder Troctomorpha, which includes all psocodean HD families (Liposcelididae and Phthiraptera), had significantly (t= 8.2, p<0.001) more aa substitutions (72) than the two DD suborders, Trogiomorpha (47) and Psocomorpha (40) (**Figure 3**, **Figure S8**, **Table S8**).

#### Diptera

The Cecidomyiidae, the only exclusively HD family of Diptera, had far more substitutions than the other 81 DD families (54 vs 34). A second dipteran family, Sciaridae, which includes some HD species, showed fewer substitutions (39) than the Cecidomyiidae, but ranked 5^th^ after Tephritidae (45), Ditomyiidae (43), and Pipunculidae (41) (**Figure S2**, **Table S3**).

#### Coleoptera

The 82 DD families of Coleoptera had an average of 36 substitutions, but two families were in the upper quintile (Phengodidae**–**57, Anamorphidae**–**53). With 37 substitutions, the sole species in the only HD coleopteran family (Micromalthidae) did not show elevated aa substitution (**Figure 3**, **Figure S1**). The only other reports of HD in Coleoptera involve two tribes in the subfamily Scolytinae, members of the hyper-diverse family Curculionidae. Among the 16 subfamilies of curculionids in our analysis, the Scolytinae had the highest mean substitution count (46) while the highest number of substitutions (50) was in the Xyloborini, a tribe which is HD (**Table S9**).

### Amino acid variation in DD insect orders

The 428 DD families exhibited significantly fewer aa substitutions than the 85 HD families (t = 14, p<0.001) (**Table S10**). The Strepsiptera was a great exception as its component families all showed many aa substitutions (x = 89) with a range from 81 (Xenidae) to 93 (Corioxenidae, Elenchidae) (**Figure 3**, **Figure S10G**, **Table S3**). All families within the Phasmatodea and Embioptera also exhibited substitutions in the upper quintile, while those in the suborder Annulipalpia of Trichoptera also crossed this threshold. (**Figure 3**, **Figure S9F**, **Figure S9G Figure S10F**).

### Indels in COI

Indels were only detected in 7 of the 428 DD families and all were in Strepsiptera. By contrast, 27 of the 85 HD families possessed indels including taxa in both HD orders (Hymenoptera, Thysanoptera) and in two of the four orders (Hemiptera, Psocodea) with some HD lineages. In total, 38 indels were detected and these included five insertions and 33 deletions (**Figure 4**, **Table S11**).

**Figure 4:**
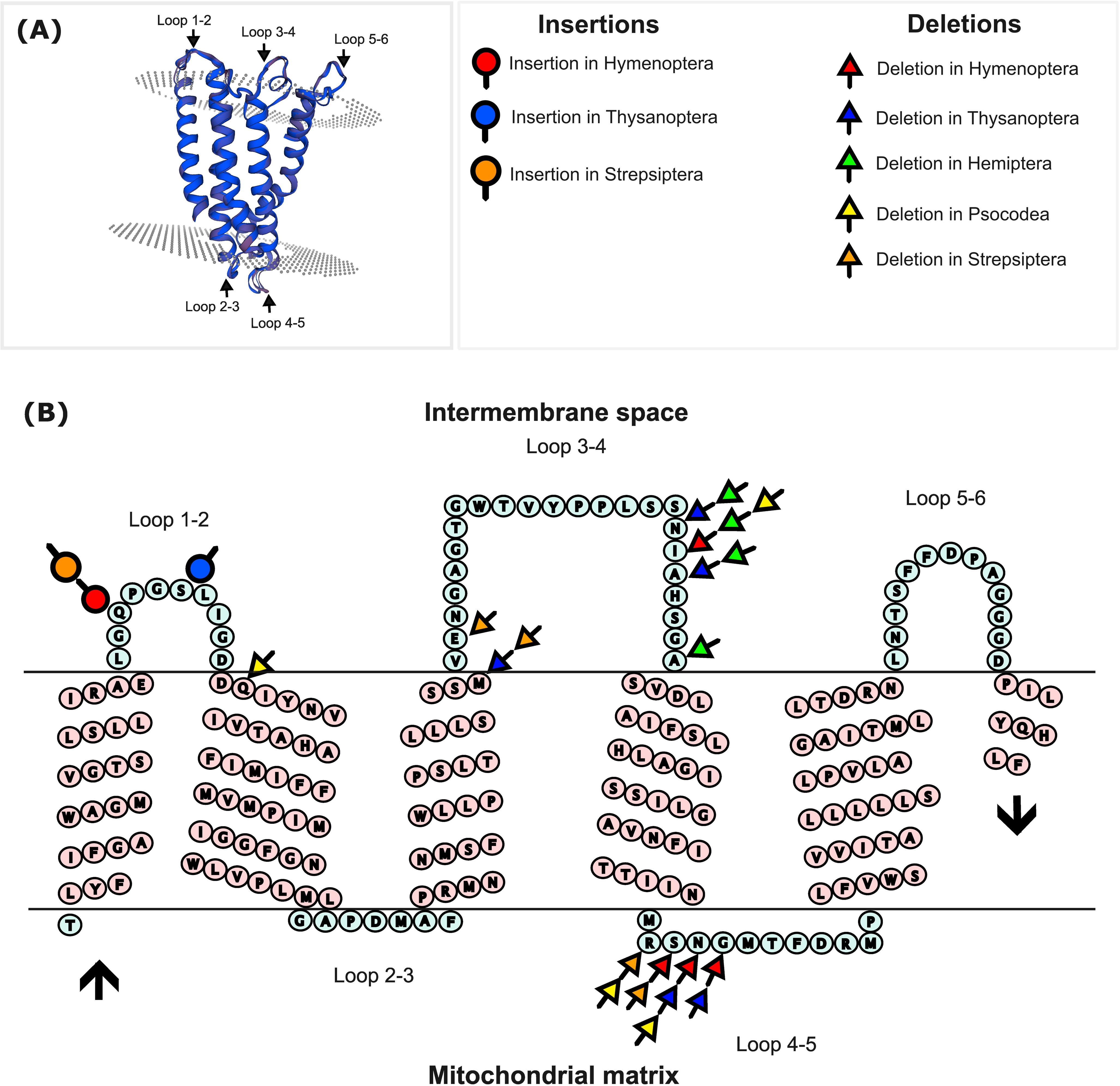
(A) Three-dimensional model of the secondary structure of the protein encoded for by COI barcode region. (B) Secondary structure of the consensus amino acid sequence for COI in insects showing the positioning of insertions and /deletion in five insect orders. The black arrows mark the N- and the C-terminal of COI

#### Insertions

Insertions were observed in three orders (Hymenoptera, Strepsiptera, Thysanoptera) and all were positioned within mitochondrial intermembrane loops 1**–**2 (**Figure 4**, **Table S11**).

#### Deletions

Deletions were both more prevalent and variably positioned than insertions. Deletions in intermembrane loops 1**–**2 were only detected in the Psocodea while deletions in intermembrane loops 3**–**4 were only observed in Hemiptera. In contrast to these deletions restricted to one order, four orders (Hymenoptera, Psocodea, Strepsiptera, Thysanoptera) possessed deletions in mitochondrial matrix loops 4**–**5 (**Figure 4**, **Table S11**).

## Discussion

The cytochrome *c* oxidase 1 protein is critical for oxidative phosphorylation, but its functionality depends upon interactions with the other two COI proteins encoded by the mitogenome plus an array of proteins encoded by the nucleus (Bar-Yaacov et al., 2012, Cunatova et al., 2020, McDiarmid et al., 2023). Although COI has the slowest evolutionary rate of the 12 protein-coding genes in the animal mitogenome (Dong et al., 2021), this study revealed amino acid substitutions at 177 of the 220 sites in its barcode region. Comparison of amino sequences across 513 insect families revealed wide variation in the incidence of substitutions among these lineages. Some taxa showing elevated substitution possess life history traits previously associated with rate acceleration such as the loss of flight (Mitterboeck and Adamowicz 2013) or a parasitic lifestyle (e.g., McMahon et al., 2011; Sweet et al., 2022). However, our analysis revealed that haplodiploid lineages possessed 1.7x as many substitutions as their diplo-diploid counterparts suggesting that this breeding system accelerates mitogenome evolution, reinforcing the evidence provided by Li et al (2024) in a genome wide scan in five HD lineages of arthropods.

Our analysis was taxonomically comprehensive as it examined all five major lineages of insects and 26 of their 29 orders (Apterygota**–**2, Palaeoptera**–**2, Polyneoptera**–**8, Paraneoptera**–**3, Holometabola**–**11). The major lineages showed two-fold variation in their number of amino acid substitutions from outgroup taxa, but the patterning of this variation did not match expectations if flight loss or parasitic lifestyle were the primary drivers. For example, all families of Apterygota (4) and Palaeoptera (22) had similar substitution counts (x = 32.5, x =32.6 respectively) although the former are flightless while the latter fly.

Substitution patterns in the Polynoptera supported the importance of flight loss in accelerating substitution rates as the 39 families in the six orders (Blattodea, Dermaptera, Mantodea, Orthoptera, Plecoptera, Zoraptera) capable of flight had fewer substitutions (x = 35.3) than the nine families of Embioptera and Phasmatodea with flightless females (x = 54.4). The three orders (Hemiptera, Psocodea, Thysanoptera) of Paraneoptera showed marked variation among lineages. For example, the three suborders (Heteroptera, Auchenorrhyncha, Sternorrhyncha) and 81 families of Hemiptera had substitution counts ranging from ranging from 31 (Corixidae) to 120 (Ortheziidae). Among the hemipteran suborders, the 38 heteropteran families, had the fewest substitutions (x = 39.8), while the 24 families of Auchenorrhyncha held middle place (x = 52.6), and the 19 sternorrhynch families had the most (x = 89.2). Among the Psocodea, the 16 free-living, winged families of Psocomorpha and Trogiomorpha all had low substitution counts (x = 40.4) while the six flightless families (five also parasitic) were high (x = 72.3). The four families in the final Paraneopteran order, Thysanoptera, had many substitutions (x = 66.2) although they are both free-living and capable of flight. The Holometabola includes 11 orders, four species-rich and seven species-poor. Most families in three megadiverse orders (Coleoptera, Diptera, Lepidoptera) had few substitutions. None of the 59 lepidopteran families and just 1 of 83 dipteran families (Cecidomyiidae) reached the upper quintile, reflecting the low mean substitution counts for families in these orders (x = 33.9, x = 37.9). The Coleoptera also had few substitutions (x = 36.0) although 4 of its 83 families were in the upper quintile and the top-ranked Phengodidae has flightless (larviform) females. By contrast, 57 of 62 families of Hymenoptera (x = 61.9) were in the upper quintile although many are free-living and capable of flight. Among the species-poor orders of Holometabola, five of seven had low levels of amino acid substitutions. Four (Mecoptera–31, Megaloptera–27, Neuroptera–31, Rhapidioptera–29) are free-living and capable of flight, but the Siphonaptera (32) are flightless and parasitic. By contrast, all seven families of Strepsiptera had many amino acid substitutions (x = 88.7), perhaps linked to their endoparasitic lifestyle and flightless females. Finally, the Trichoptera showed a divide with 16 families having low amino acid substitution (x= 43) while six other families were in the high quintile (x = 58). All trichopterans are free-living but the families with the higher substitution counts are net spinners which that are less mobile than the other families.

This study has revealed complex patterns of variation in the levels of amino acid substitution in the COI barcode region among insect lineages. Some taxa lacking flight (e.g., Apterygota) or even combining it with a parasitic lifestyle (e.g., Siphonaptera) showed few substitutions while other lineages capable of flight and free-living, such as Thysanoptera, had many substitutions. These cases suggest that factors beyond flight loss and a parasitic lifestyle modulate the pace of mitogenome evolution and this study identifies haplodiploidy as a key candidate. Nearly all families in the two HD orders (Hymenoptera, Thysanoptera) were placed in the upper quintile. In the five other orders that include both HD and DD lineages, the former taxa show more amino acid substitutions. Trait covariation makes it difficult to disentangle the relative impacts of flight loss, parasitism, and breeding system in accelerating the pace of mitogenome evolution. For example, the hemipteran families with the highest number of amino acid substitution are haplodiploid, but their females are also flightless and plant parasites. Likewise, the rapid evolutionary rate observed in the Phthirapteran lineage within Psocodea can be ascribed to the synergistic effect of all three factors. Despite such covariation, the important role of haplodiploidy in accelerating mitogenome evolution is suggested by cases where key variables known to influence mitogenome evolution vary within an order. For example, 21 families of Diptera are parasitic (Eggleton and Belshaw, 1992), but only one, the Cecidomyiidae was in the upper quintile, and it is the only dipteran family known to be exclusively haplodiploid. Moreover, among of the 16 curculionid subfamilies analyzed, the Scolytinae had the highest mean substitution rate and the sole HD tribe in our study, the Xyloborini, had the highest individual substitution rate.

While amino acid substitutions in COI are likely to impact fitness, indels are even more likely to have selective impacts (Lin et al., 2017). Prior studies have shown that indels in exonic regions are short, typically 1-2 codons in length, with deletions about twice as common as insertions (Chaux et al., 2007), patterns similar to those noted in this study. Aside from revealing that indels were restricted to particular segments of COI, they were only present in HD taxa with the exception of several cases in the Strepsiptera, an order showing an exceptional level of amino acid substitution. This congruence suggests that factors favouring high rates of amino acid substitution also facilitate indels. The traits suggest the value of careful studies to confirm that strepsipterans are indeed DD. Although cytological studies have not indicated that their males are haploid, cases of haplodiploidy involving chromosome inactivation are easily overlooked. Known from six insect orders (Blackmon et al., 2017) and nine arthropod lineages (Wang & Davis, 2014), it is worth confirming that it does not also occur in Strepsiptera.

Although this study has only examined patterns of amino acid divergence in a segment of one of the 12 protein-coding genes in the mitogenome, prior studies have established that this region is a strong proxy for the mitogenome (Hebert et al., 2016; Roslin et al., 2022). As such, elevated rates of non-synonymous substitutions and indels are likely to characterize the mitogenomes of haplodiploid species, a prediction supported by the genomic studies undertaken by Lin et al (2024).

What accounts for the linkage between haplodiploidy and elevated amino acid substitutions? Prior studies have identified strong coevolution between mitochondrial and nuclear gene products. Mitonuclear compensatory coevolution is thought to play a strong role in evolutionary dynamics (Li et al., 2017; Yan et al., 2019; Piccinini et al., 2021) and the impact of this coadaptation on gene flow and speciation has received recent attention (Hill, 2020. Because of this coevolution, many, if not all, newly arisen mitochondrial mutations which increase fitness will require compensatory substitutions in nuclear genes. Because most variation in the nuclear genome is recessive or weakly penetrant, it will not be exposed to selection in DD taxa. However, such genes gain full expression in males of haplodiploids. Although they do not transmit mitochondria, these males can transmit nuclear genes with favorable impacts on mitochondrial functioning. By exposing genes with favorable effects on nuclear-mitochondrial coevolution to selection, haplodiploidy facilitates the fixation of newly arisen mitochondrial mutations that would otherwise be lost to drift or rejected by selection. The pervasive impacts of Muller’s ratchet (Muller, 1964) may drive mitochondrial gene rearrangements (Moreno-Carmona et al., 2021) and even the fragmentation of mitogenomes (Chinnery et al., 2019; Sweet et al., 2022) in HDs. Future studies which examine whole genomes will undoubtedly advance our understanding of the mitonuclear interactions and the impacts of breeding systems on them. However, future investigations should also test the generality of patterns revealed by the analysis of insects by considering more haplodiploid lineages within the Arthropoda (e.g., mites) and other phyla (e.g., Nematoda, Rotifera). If these investigations confirm that haplodiploidy is an evolutionary accelerant, work should also compare organellar evolution in groups whose life cycles are largely haploid (e.g., mosses) with those which are diploid (e.g., higher plants, ferns). Protistans would seem particularly fertile ground for such investigations as their life cycles span the spectrum from haplontic to diplontic (Mable and Otto, 1998).

## Ethics

This study did not require ethics approval by a human subject or animal welfare committee.

## Data accessibility

All data in this article are available on BOLD (www.boldsystems.org) in the dataset: DS-HDMS at doi: dx.doi.org/10.5883/DS-HDMS.

## Declaration of AI use

AI was not used to create this manuscript.

## Authors’ contributions

AP: data analysis, first draft; PDN: conceptualization, writing-review, and editing.

## Acknowledgements

We thank all researchers who have contributed public records to BOLD.

## Figure captions

**Figure S1:** Radar plot of the number of amino acid substitutions in the barcode region of COI between a dipluran outgroup and 83 families of Coleoptera. The blue annulus indicates 50 substitutions.

**Figure S2:** Radar plot of the number of amino acid substitutions between a dipluran outgroup and 83 families of Diptera. The blue annulus indicates 50 substitutions.

**Figure S3:** Radar plot of the number of amino acid substitutions between a dipluran outgroup and 59 families of Lepidoptera. The blue annulus indicates 50 substitutions.

**Figure S4:** Radar plot of the number of amino acid substitutions between a dipluran outgroup and 62 families of Hymenoptera. The blue annulus indicates 50 substitutions.

**Figure S5**: Radar plot of the number of amino acid substitutions between a dipluran outgroup and 81 families of Hemiptera. The blue annulus indicates 50 substitutions

**Figure S6:** Radar plot of the number of amino acid substitutions between a dipluran outgroup and two orders of Apterygota: (A) 2 families of Archeognatha and (B) 2 families of Zygentoma. The blue annulus indicates 50 substitutions.

**Figure S7:** Radar plot of the number of amino acid substitutions between a dipluran outgroup and two orders of Palaeoptera: (A) 11 families of Ephemeroptera, and (B) 11 families of Odonata. The blue annulus indicates 50 substitutions.

**Figure S8:** Radar plot of the number of amino acid substitutions between a dipluran outgroup and two orders of Paraneoptera: (A) 22 families of Psocodea and (B) 4 families of Thysanoptera. The blue annulus indicates 50 substitutions.

**Figure S9:** Radar plot of the number of amino acid substitutions between a dipluran outgroup and 7 orders of Polyneoptera: (A) 4 families of Dermaptera, (B) 9 families of Plecoptera, (C) 12 families of Orthoptera, (D) 4 families of Mantodae, (E) 9 families of Blattodea, (F) 6 families of Phasmatodea, and (G) 3 families of Embioptera, (H) 1 family of Zoraptera

**Figure S10:** Radar plot of the number of amino acid substitutions between a dipluran outgroup and six orders of Holometabola: (A) 2 families of Raphidioptera, (B) 2 families of Megaloptera, (C) 5 families of Neuroptera, (D) 4 families of Siphonaptera, (E) 3 families of Mecoptera, (F) 22 families of Trichoptera, and (G) 7 families of Strepsiptera. The blue annulus indicates 50 substitutions.

## Table captions

**Table S1**: Insect orders, families, and number of barcodes in the dataset.

**Table S2:** Regression of the number of amino acid substitutions between the consensus COI sequences for Diplura and Protura and the consensus COI sequence for each of 513 insect families.

**Table S3:** Number of amino acid substitutions between the dipluran outgroup and each of the 513 insect families in 26 orders.

**Table S4:** Mean number of amino acid substitutions and ANOVA of the five lineages (Paraneoptera, Holometabola, Polyneoptera, Palaeoptera, Apterygota) in the class Insecta.

**Table S5:** Mean number of amino acid substitutions and ANOVA examining differences among the five major insect orders (Coleoptera, Diptera, Hemiptera, Hymenoptera, Lepidoptera).

**Table S6:** Amino acid substitutions and statistics for the order Hymenoptera.

**Table S7:** Amino acid substitutions and statistics for the order Hemiptera.

**Table S8:** Amino acid substitutions and statistics for the order Psocodea.

**Table S9:** Amino acid substitutions in subfamilies of the Curculionidae (Coleoptera).

**Table S10:** t test of amino acid substitutions between all HD and DD families.

**Table S11:** Location of insertions and deletions in the secondary structure of the consensus amino acid sequence for COI.

